# The effective population size and mutation rate of influenza A virus in acutely infected individuals

**DOI:** 10.1101/2020.10.24.353748

**Authors:** John T. McCrone, Robert J. Woods, Arnold S. Monto, Emily T. Martin, Adam S. Lauring

## Abstract

The global evolutionary dynamics of influenza viruses ultimately derive from processes that take place within and between infected individuals. Recent work suggests that within-host populations are dynamic, but an *in vivo* estimate of mutation rate and population size in naturally infected individuals remains elusive. Here we model the within-host dynamics of influenza A viruses using high depth of coverage sequence data from 200 acute infections in an outpatient, community setting. Using a Wright-Fisher model, we estimate a within-host effective population size of 32-72 and an *in vivo* mutation rate of 3.4×10^−6^ per nucleotide per generation.

## Introduction

The rapid evolution of influenza viruses places demographic processes such as population growth, transmission, and epidemiological spread on a similar time scale as the accumulation of genetic substitutions. This similarity of scale makes it possible to infer demographic processes from genetic sequence data using phylodynamic methods (Lemey et al. 2009; Bedford et al. 2014; Bedford et al. 2015). Investigations of the global dynamics of influenza have been successful, in part, because the complexities of within- and between-host processes can be collapsed into a limited number of parameters in the coalescent or birth-death process when averaged over large spatial and temporal scales. However, it becomes increasingly important to disentangle these processes to address more granular questions; for example, the transmission of viruses at local scales or selective pressures imposed by vaccines, antivirals, or novel hosts.

Phylogenetic approaches that separate within-host processes from those acting at epidemiological scales rely on simple population genetic models to capture the complex dynamics that occur within infected individuals (Didelot et al. 2014; Hall et al. 2015; Didelot et al. 2017; De Maio et al. 2018). However, the accuracy of these models depends on reliable estimates of the within-host effective population size (N_e_), which in the case of influenza virus, has proven difficult due to inherent challenges in collecting longitudinal samples from representative infections. Here, we take advantage of a large, well-studied, community cohort with robust deep sequencing data, from which two important results have emerged (McCrone et al. 2018). First, within-host selection for novel antigenic variants is weak, and second, transmission between hosts imposes a significant bottleneck on the viral population. We leverage these findings to fit a Wright-Fisher model to capture the dynamics of within-host populations. This model provides consistent and robust estimates of the within-host N_e_ and mutation rate of influenza A virus (IAV) when applied to cross-sectional and longitudinal samples. These findings provide an important baseline for defining processes related to the local dynamics of IAV, and of RNA viruses in general.

## Results

We recently performed high depth of coverage sequencing of 249 IAV populations recovered from 200 individuals enrolled in the Household Influenza Vaccine Effectiveness (HIVE) study (McCrone et al. 2018). This large number of samples collected within a prospective community-based cohort is a rich dataset for exploring influenza virus evolution over the course of a natural infection. In this and other works, we have documented our sensitivity and specificity for detection of intrahost single nucleotide variants (iSNV) and our error in allele frequency measurement (McCrone and Lauring 2016; Debbink et al. 2017; McCrone et al. 2018). Our dataset also includes 49 serially sampled individuals, who provided a self-collected specimen at the time of symptom onset and a clinic-collected specimen 0–7 days later. This affords an opportunity to explore changes in viral populations in naturally infected individuals over a short time scale.

We applied a continuous diffusion approximation of the Wright-Fisher model to define the within-host accumulation of mutations using 196 cross-sectional samples, collected 1-7 days following the onset of symptoms (Rouzine et al. 2001). Because we have previously estimated an effective transmission bottleneck of 1-2 genetically distinct variants, we made the simplifying assumption that each infection was clonal and modeled the accumulation of diversity until the time of sampling as a neutral process. Maximum likelihood optimization of this model estimated an *in vivo* mutation rate of 3.4×10^−6^ (95% CI 3.1-3.7×10^−6^) mutations per nucleotide per generation (6 hours) and a within-host N_e_ of 36 (95% CI 31-41, Figure 1). We have recently estimated that the majority of mutations in IAV are detrimental and therefore unlikely to be observed at detectable frequencies (Visher et al. 2016). As only ~10% of mutations in influenza A virus are neutral, we propose that the true *in vivo* mutation rate is approximately ten-fold higher than our estimated rate, which does not account for purifying selection. This results in an *in vivo* mutation rate of approximately 3.4 × 10^−5^ substitutions per nucleotide replicated per generation, which is within the range of estimates for IAV’s biochemical mutation rate in epithelial cells (Sanjuán et al. 2010).

**Figure 1.**
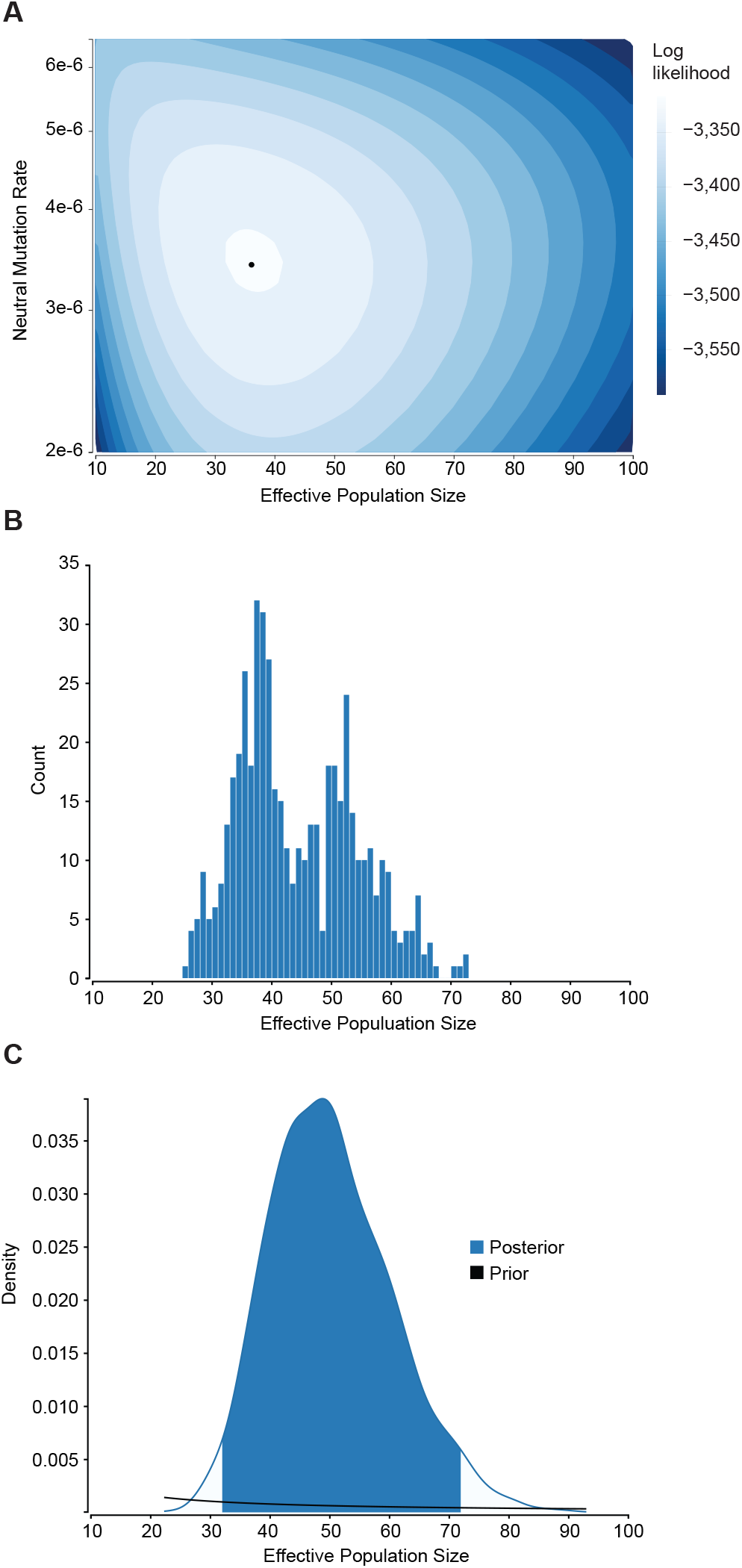
(A) Joint estimate of within-host mutation rate and effective population size. Contour plot shows the log likelihood surface for estimates of the effective population size and neutral mutation rate. The point represents the peak (μ = 3.4×10^−6^, N_e_ = 36). Log likelihoods for each contour are indicated. (B) The distribution of N_e_ estimated in 500 subsamples of the data in which one iSNV was taken per individual. The bimodality of the distribution reflects a slight sensitivity to the inclusion of a few specific iSNV. (C) The posterior and prior probability densities for N_e_ over all values explored in the in the combined MCMC chains (22-93). The 95% HPD of the posterior (32-72) is shaded blue.

To determine the robustness of our N_e_ estimate, we fit this same model to changes in allele frequencies observed in a subset of paired longitudinal samples. We restricted this analysis to alleles observed at the first time point in samples taken at least 1 day apart (63 iSNV in 29 sample pairs). There was very little change in iSNV frequency in populations sampled twice on the same day (R^2^ = 0.986, Figure 2, Supplement 1A of (McCrone et al. 2018)). The concordance of same-day samples suggests that our sampling procedure and frequency measurements are reproducible. Maximum likelihood optimization of this model revealed a within-host N_e_ of 34 (95% CI 25-46, Table 1), very similar to that observed in the cross-sectional data above. Comparable estimates were obtained when synonymous and nonsynomous mutations were fit separately (Table 1). As there is some uncertainty in the within-host generation time (Geoghegan et al. 2016), we also estimated the N_e_ based on a 12 hour generation. As expected, increasing the generation time results in a smaller N_e_.

**Table 1.** Within host effective population size of IAV.

| iSNV Used | Generation Time (h) | Effective Population Size (95% CI) |
| --- | --- | --- |
| All | 6 | 34 (25-46) |
| All | 12 | 17 (13-23) |
| Nonsynonymous | 6 | 27 (16-44) |
| Synonymous | 6 | 40 (27-59) |
| All | 12 | 17 (13-23) |
| Nonsynonymous | 12 | 14 (8-22) |
| Synonymous | 12 | 20 (14-29) |

The Wright-Fisher model assumes that each allele in a population is independent. This assumption would be violated if there were multiple iSNV per genomic segment or varying linkage of iSNV across segments due to reassortment. However, heterotypic reassortment is quite rare within hosts (Sobel Leonard et al. 2017), and the per-sample diversity in our dataset was sufficiently low that nearly all segments had either 0 or 1 iSNV. To ensure that our results were robust to the assumption of independent allele frequencies, we fit the above model 500 times, each time randomly subsetting our data such that only one iSNV per individual was included. In practice, this approach also tested the sensitivity of our estimates to individual allele trajectories. Under these conditions, we found a median N_e_ of 42 (IQR 37-52, Figure 1B). Thus, in the initial analysis, non-independence among iSNV within the same host may have caused a slight bias due to a few hosts with extreme frequency changes.

The estimates above include the probability that undetected variants are present but missed due to imperfect sensitivity (see Methods and (McCrone and Lauring 2016)); however, they do not account for uncertainty in the frequency measurements, which if large, would bias the N_e_ estimate toward lower values. To accommodate this uncertainty we relied on the fact that 141 of the 249 samples in were amplified and sequenced in duplicate (McCrone et al. 2018). We modeled the frequency-dependent variance present in the data as a beta distribution with *α* = *p* ∗ *n*, *β* = *p*(1 − *p*) ∗ *n*, where *p* represents the true frequency (the mean in the duplicate measurements) and *n* roughly represents the number of samples in a binomial distribution with probability *p*, and was determined with maximum likelihood optimization. We then adapted a Bayesian approach and estimated the posterior distribution of N_e_ integrated over all unobserved, true frequency trajectories. The analysis resulted in a marginally increased N_e_ estimate of 50 (32-72 95% HPD, Figure 1C). The agreement between this model and our previous estimates suggests that the relatively small N_e_ is driven by the allele trajectories themselves and is not the result of uncertainty in our frequency measurements.

## Discussion

We have investigated the within-host dynamics of influenza in a large, well-defined cohort of representative infections and found that, under a Wright-Fisher model, the population is characterized by a small effective population size. Our findings differ from those reported in studies of immunosuppressed, chronically infected individuals, which have shown that within-host populations of influenza virus are characterized by large effective population sizes, clonal interference, and selective pressures that mimic those seen at larger biological scales (Xue et al. 2017; Lumby et al. 2020). The difference in these N_e_ estimates likely lies in the fundamental difference between the population dynamics of acute and chronic infections. Chronic infections, which manifest in rare immunologically atypical hosts, establish large, stable populations and may be “insulated” from the drastic fluctuation in population size that define acute cases. In the absence of any evidence for antigenic selection, it seems that evolution during the early period of influenza infections, the time frame during which transmission is most likely to occur, is best modelled as a stochastic process.

The Wright-Fisher model provides a simple framework for exploring the evolutionary dynamics of “real-world” populations. The model’s tractability comes at the cost of many simplifying assumptions (e.g. constant population size, discrete generations, homogenous mixing, neutral evolution), which are rarely, if ever, met by biological populations. Influenza viruses clearly exist as complex populations whose evolutionary dynamics are influenced by a mixture of processes not captured explicitly in the Wright-Fisher model (e.g. deleterious mutation load, migration between sites of infection, rapid population growth and decline (Lakdawala et al. 2015; Visher et al. 2016; Zhao et al. 2019)). However, the detailed, longitudinal sampling needed to fit models that explicitly capture this complexity is not available for most influenza infections, which are typically short-lived and not medically attended. In the absence of such data, we have chosen a more tractable model that can yield reliable estimates of the general tendencies, rather than more complex models that may lack identifiability and generalizability.

These estimates of the effective population size and mutation rate, combined with previous estimates of the transmission bottleneck, provide a useful expectation for the shared diversity between direct transmission pairs, and can be used in conjunction with standard epidemiological models to study the forces that drive influenza evolution at a granular level.

## Methods

### Fitting mutation rate and N_e_

The diffusion approximation to the Wright-Fisher model makes predictions regarding the allele frequency spectrum of a population given a mutation rate and N_e_. Starting from a monomorphic state, while t≪N_e_, the probability of observing a mutation at frequency *p*_*t*_ be approximated as in equation 85 of (Rouzine et al. 2001)

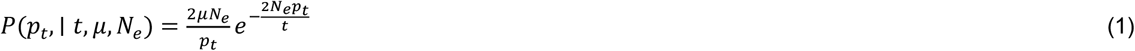

Where *μ* is the mutation rate in substitutions/site/generation, N_e_ is the effective population size and *t* is the number of generations. Consistent with previous models of within-host influenza, we set the generation time to 6 hours (Geoghegan et al. 2016). We further assumed that infection began 1 day prior to symptom onset (Carrat et al. 2008).

To account for limitations in iSNV detection, we integrated over regions of the probability density where we have observed less than perfect sensitivity. The probability of not observing an iSNV at a locus is given by summing over the possibilities that (i) a mutation is present but below our level of detection *P*(*p_t_* ≈ 0 | *p*_*t*_ < 0.02, *t*, *μ*, *N*_*e*_), and (ii) a mutation is present but missed due to low sensitivity at low frequencies *P*(*p*_*t*_ ≈ 0 | 0.02 < *p*_*t*_ < 0.1, *t*, *μ*, *N*_*e*_). In this model, we assumed there were 13,133 polymorphic loci in each sample (the number of coding sites present in the reference strain from 2014-2015). Under these assumptions,

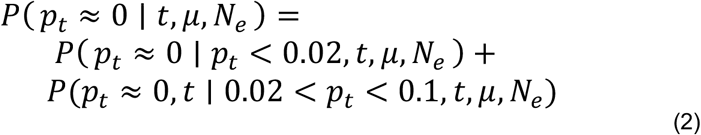

Where

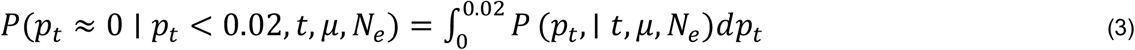

and

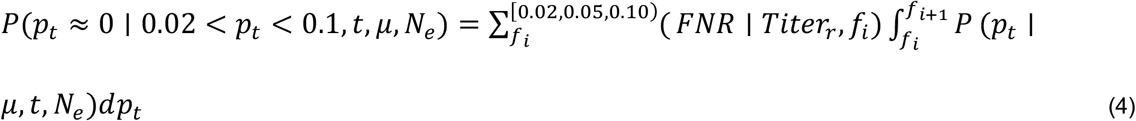

Where (*FNR* | *Titer*_*r*_, *f*_*i*_) is the false negative rate given the frequency and the sample titer (See Supplementary File 1 in (McCrone et al. 2018)). As before, we assumed the sensitivity in the intervals between 0.02, 0.05 and 0.1 was equal to the sensitivity at the lower bound, and that the sensitivity was perfect at frequencies above 0.1. The log-likelihood of a given *μ* and *N*_*e*_ pair is then the sum of the log of equations 1 and 2 for all possible sites in the data set. The maximum-likelihood values were estimated using the bbmle package in R (Ben Bolker and Team 2020; Team 2020).

### Diffusion approximation

We implemented the diffusion approximation as in (Kimura 1955), with minor modifications. As above, we included the limitations in our sensitivity to detect rare iSNV by integrating over all possible explanations for why an iSNV might not be observed at the second time point.

### Bayesian implementation of the diffusion approximation

To account for measurement error in our estimates we adopted a similar approach to that developed in (Williamson and Slatkin 1999). The likelihood of observing frequencies 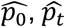 at time 0 and t given the true frequencies *p_0_* and *p*_*t*_

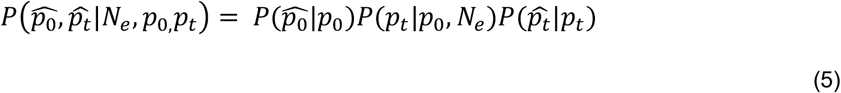

where 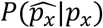 counts for measurement error and is defined for 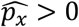 by the probability density at 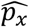 of a beta distribution with *α* = *p* ∗ *n*, *β* = *p*(1 − *p*) ∗ *n* where n=503 and was determined from the estimating the error in replicate sequencing samples.

In cases where 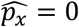 and *p*_*x*_ > 0, 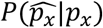 is the sum of the cumulative density function of the same beta distribution up to 0.02 (i.e. the variant is detected below the limit of detection) and the probability of not detecting the variant given the sample titer and false negative rate as above (the variant was not observed to imperfect sensitivity). *P*(*p*_*t*_|*p*_0_, *N*_*e*_) is the transition probability of a variant at frequency *p*_0_ to drifting to *p*_*t*_ given t generations and an the effective population size of *N_e_* as in equation 15’ in (Kimura 1955). The posterior is proportional to the product of this likelihood and priors on N_e_, p_0_, and p_t_. We choose uniform priors for *p_0_* and *p*_*t*_ and a diffuse gamma prior with shape of 0.036 and scale of 1000 (mean 36 as informed by the cross-sectional data analysis). As with the other analyses the generation time was set to 6 hours (Geoghegan et al. 2016). This approach was implemented as a plugin for BEAST and the posterior was estimated using BEAST v1.10.4 (Suchard et al. 2018). Ten independent MCMC chains were run for 10 million states. Each chain was sampled every 10,000 iterations with the first 1 million states discarded as burn in. All ten chains were combined and ESS for all parameters was >200. Convergence was assessed in *TRACER* (Rambaut et al. 2018).

## Acknowledgments

This work was supported by a Clinician Scientist Development Award from the Doris Duke Charitable Foundation (CSDA 2013105) and R01 AI118886 to ASL. The HIVE cohort was supported by NIH R01 AI097150 and CDC U01 IP00474 to ASM. JTM was supported by the Michigan Predoctoral Training Program in Genetics (T32GM007544). RJW was supported by K08AI119182. We thank Alexey Kondrashov and Aaron King for helpful discussion.

## Data availability

All raw sequence data have been deposited at the NCBI sequence read archive (BioProject Accession number PRJNA412631) as described in (McCrone et al. 2018). Variants were called following the validated protocol outlined in (McCrone and Lauring 2016) with details provided in (McCrone et al. 2018). Called variants and the scripts needed to reproduce this analysis are publicly available at https://github.com/lauringlab/IAV_within-host_Ne

